# Development of equations for converting random-zero to automated oscillometric blood pressure values

**DOI:** 10.1101/688895

**Authors:** Li Yan, Xiaoxiao Wen, Alan R Dyer, Haiyan Chen, Long Zhou, Paul Elliott, Yangfeng Wu, Queenie Chan, Liancheng Zhao

## Abstract

**Objective:** This study aimed to collect data to compare blood pressure values between random-zero sphygmomanometers and automated oscillometric devices and generate equations to convert blood pressure values from one device to the other.

**Methods:** Omron HEM-907, a widely used automated oscillometric device in many epidemiologic surveys and cohort studies, was compared here with random-zero sphygmomanometers. Two hundred and one participants aged 40-79 years (37% men) were enrolled and randomly assigned to one of two groups with blood pressure measurement first taken by automated oscillometric devices or by random-zero sphygmomanometers. The study design enabled comparisons of blood pressure values between random-zero sphygmomanometers and two modes of this automated oscillometric device – automated and manual, and assessment of effects of measurement order on blood pressure values.

**Results:** Among all participants, mean blood pressure levels were lowest when measured with random-zero sphygmomanometers compared with both modes of automated oscillometric devices. Several variables, including age and gender, were found to contribute to the blood pressure differences between random-zero sphygmomanometers and automated oscillometric devices. Equations were developed using multiple linear regression after taking those variables into account to convert blood pressure values by random-zero sphygmomanometers to automated oscillometric devices.

**Conclusion:** Equations developed in this study could be used to compare blood pressure values between epidemiologic and clinical studies or identify shift of blood pressure distribution over time using different devices for blood pressure measurements.

## Introduction

Due to the high prevalence of high blood pressure (BP) worldwide and its strong association with increased risk for cardiovascular disease (CVD), BP measurement is standard in clinical practice. In order to provide an accurate estimate of BP level and assess its influence on cardiovascular health, correct procedures and acceptable precision of the devices used for BP measurement are crucial in clinical and epidemiologic studies.

The standard mercury sphygmomanometer (HgS), with a stethoscope to detect Korotkoff sounds [1], has been recognized as the gold standard for measuring BP for many years, though biases in this procedure, e.g. observer bias due to digit preference, may cause inaccurate measurements [2]. The random-zero sphygmomanometer (RZS) was introduced later as a modified device to measure BP, for the purpose of reducing observer bias. Many large epidemiologic studies until the late 1990s used RZS to measure BP, including the International Study on Macronutrients and Blood Pressure (INTERMAP) [3].

However, a number of studies have cast doubt on the accuracy of the RZS, and found that it underestimates BP level compared with HgS. Brown et al. [4] reported that random-zero values may not be randomly distributed, e.g. when speedy measurement is carried out without sufficient cuff inflation for filling of the diaphragm chamber. The European Society of Hypertension (ESH) did not recommend using the RZS in clinical practice because it failed the quality standards of the Association for the Advancement of Medical Instrumentation (AAMI) and the British Hypertension Society (BHS) [5]. Another concern is the threat from mercury to human health and the environment. The United States and the European Union decided in 1999 to phase out the use of mercury instruments and use non-mercury devices instead [6]. Thus, the RZS (or HgS) cannot be used in longitudinal studies anymore, including the INTERMAP China Prospective (ICP) Study, and an alternative non-mercury device must be used to measure BP in these studies.

Most recent studies have used automated oscillometric devices (AODs) for BP measurements; these detect pulse wave oscillations and estimate BP values via algorithms. Protocols were developed to validate these devices before recommendations for use [5, 7].

At the individual level, long-term BP change has been shown to predict CVD risk [8-10], which can also be used as a primary outcome in prospective studies. At the population level, consistent information is needed to understand elevated BP and its temporal trend and how the BP distribution has shifted over time. Investigation of temporal trends of BP would be hampered if different devices for BP measurement are used, since systematic measurement differences exist among devices. In order to make the measurements by different devices comparable, equations need to be developed to inter-convert BP values. Here, we report results from a BP calibration study designed to collect data for converting RZS to Omron AOD BP values.

## Methods

### Study sample and participants

Three hundred twenty-five persons were screened for the current study and 201 eligible participants aged 40-79 years were enrolled from a rural village in Beijing. The study had ethics approval from the Research Ethics Committee of Fu Wai Hospital, Beijing (reference number 2015-650) and written informed consent was obtained from each participant at the beginning of the study. A screening BP test was carried out by an Omron AOD (SINGLE Mode) at the beginning of recruitment. Participants were then enrolled into one of three SBP ranges: low <120 mm Hg (N=49); medium 120-159 mm Hg (N=119); high ≥160 mm Hg (N=33), to ensure the parameter estimate was stable and the equation was validated across all BP levels. Normal sinus rhythm was required for all eligible participants of this study, as heart arrhythmia may lead to incorrect BP readings of AODs. Persons with irregular heartbeats detected and recorded by the AODs at screening were not included in the study, i.e., those having a heartbeat rhythm that varied by more than 25% from the average heartbeat detected during BP measurement. Persons with a right arm circumference exceeding 42 centimetres and requiring use of a thigh cuff or persons whose BP of right arm cannot be measured were also ineligible.

### Comparison of automated oscillometric and random-zero devices

The Omron HEM-907 is a digital brachial BP monitor which is suitable for use in clinical settings and has been validated by standards of ESH [11], BHS [12, 13] and AAMI [13, 14]. It can measure BP both automatically (SINGLE and AVG. Modes) and manually (MANU. Mode), by combining oscillometric and auscultatory use in one model. The automatic oscillometric measurements of Omron HEM-907 are based on smart inflate technology (IntelliSense Omron), where the inflation is by a pumping system and the deflation is by a pressure-releasing electromagnetic control valve that allows rapid air release [14]. The manual mode of this monitor enabled BP auscultatory measurement with a stethoscope and manual deflation control. It has been used in several large epidemiologic studies in recent years, including The Coronary Artery Risk Development in Young Adults (CARDIA) [15] Study, the National Health and Nutrition Examination Survey (NHANES) [16], the Health Survey for England [17], and the Systolic Blood Pressure Intervention Trial (SPRINT) [18]. A Hawksley RZS (US model) [19] and the Omron HEM-907 were used in the current study. Both devices were regularly maintained by equipment professionals prior to and during the calibration study.

### Enrolment, randomization and blood pressure measurement

Every participant who entered screening completed a short questionnaire with basic information on demographics, medical history and had a screening test of BP measurement (using the Omron device in Single Mode). The BP values of eligible participants were then used to determine the target inflation pressure of the Omron device in the following procedures.

Each eligible participant was randomly assigned (stratified by range of screening SBP) to Procedure A or Procedure B for BP measurements (**Fig 1**). The randomization of measuring order was to assess its effects on BP readings as reported by previous studies [20, 21]. In Procedure A, BP measurements were first taken three times for each participant by Omron HEM-907 Single Mode (OSM), and then by RZS and Omron Manual Mode (OMM) simultaneously after 3 minutes rest. In Procedure B, participants had three BP measurements by RZS and OMM simultaneously, followed by OSM after 3 minutes rest.

**Fig 1.**
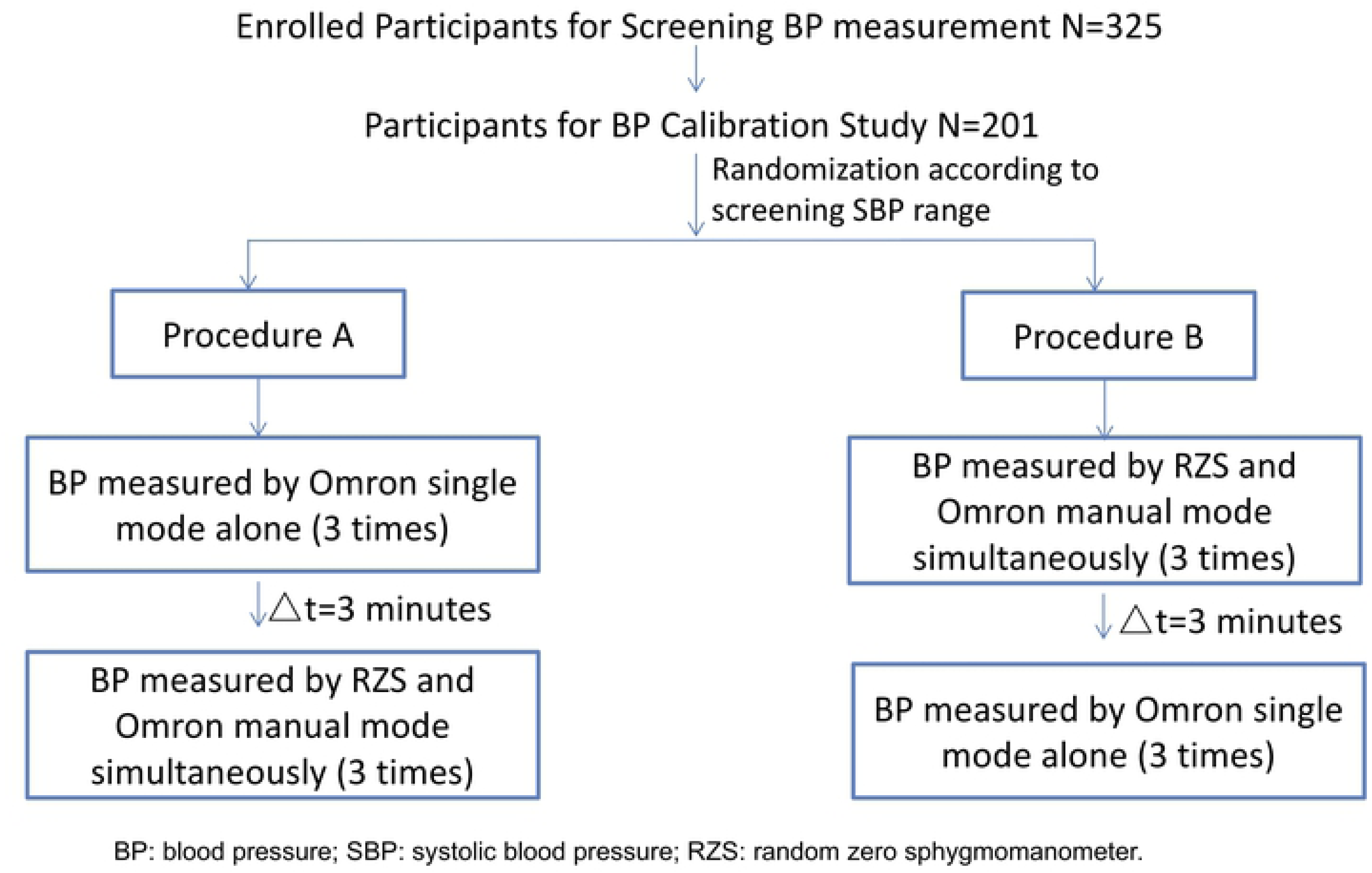
Flow chart for the blood pressure calibration study.

When BP was measured by RZS and OMM simultaneously, the two devices and an Omron cuff were connected with a “Y” tube, and the cuff was inflated by the Omron machine. “MANUAL” mode was used, and two observers listened for the Korotkoff sounds with the stethoscope to record SBP (Korotkoff I, the pressure at which a sound is first heard) and DBP (Korotkoff V, the pressure at which the sounds disappear).

Extensive efforts were adopted to reduce measurement errors between observers. The observers attended training sessions regularly and passed examinations to be certified for using both devices according to the standardized international protocols (e.g., Manual of Operations of the INTERMAP Study for RZS BP measurement). Each observer had made adequate practice of BP measurements with feedback from senior investigators before the calibration study began. During BP measurement, the two devices were positioned separately and the two observers took the measurements independently without any communication, so that each observer could not know the BP readings by the other observer. The two observers took turns using the RZS or Omron device. In addition, during data collection, a senior investigator monitored the procedures of the observers and audited them regularly to ensure that observers measured BP according to the standardized international protocols and to assure the quality of the data.

### Statistical analyses

For each BP measuring device (the RZS or the Omron HEM-907), the first measurement was discarded, as the first reading is usually systematically higher than following serial BP measurements [22]. Mean SBP and DBP were thus calculated from the second and third readings. Mean BP values were categorized according to the JNC7 classification [23].

Descriptive statistics for continuous (means and standard deviations, SDs) and categorical variables (frequencies and percentages) were calculated and are presented separately for participants by procedure (A and B) and *t*-tests for two independent samples were used to evaluate the differences between the two groups. Mean SBP and DBP by the OSM, the OMM, the RZS and the mean difference of each device/mode are presented by procedure (A and B). Paired *t*-tests were used to evaluate the differences between OSM (or OMM) and RZS BP values.

Potential factors accounting for of BP differences between the OSM and the RZS, including gender, age, device order, and their interaction terms were explored.

The difference between the two devices was plotted against the mean for the two devices according to the method developed by Bland and Altman [24], to graphically illustrate the individual differences of the BP readings by measuring devices. In addition, multiple linear regression models were used to establish the equations to convert RZS BP values to OSM BP values. Model fit was checked by using adjusted R-squared and analyses of the variance of residuals (root mean square error). As both RZS and AOD might have measurement errors, Deming Regression was used to determine if the Deming Regression equation differs from the equation obtained from multiple linear regression as sensitivity analyses (shown in **Supplementary materials**).

The observed percent agreement and kappa coefficient for BP classes (normal [<120 mm Hg], pre-hypertension [≥120 mm Hg and <140 mm Hg], and hypertension [≥140 mm Hg] according to JNC7 [23]) were calculated between the OSM and RZS, and between the OSM and calibrated RZS BP values.

All analyses were performed using SAS (Statistical Analysis System, Version 9.3 for Windows; SAS Institute Inc., Cary, North Carolina, USA). All statistical tests were two-sided, with a significance level of 0.05.

## Results

### Descriptive statistics

Comparisons of the 201 individuals (70 men and 131 women) randomized to Procedure A (n=98) or to Procedure B (n=103) are presented in **Table 1**. For participants assigned to Procedure A (mean age of 60.4 years and 37% men), BP was measured first using OSM followed by RZS/OMM measurements. For participants assigned to Procedure B (mean age of 58.8 years and 33% men), BP was measured first using RZS/OMM and then by OSM. The mean screening SBP was similar between the two groups with 136.7 mm Hg for Procedure A and 137.2 mm Hg for Procedure B. Age, gender, and screening SBP and DBP were not significantly different between the two groups.

**Table 1.** Characteristics at the screening test of the 201 participants randomized into Procedure A or Procedure B in the blood pressure calibration study.

|  | Procedure A:<br>Omron single mode<br>followed by RZS/Omron<br>manual mode (n=98) |  | Procedure B:<br>RZS/Omron manual mode<br>followed by Omron<br>single mode (n=103) |  | P value |
| --- | --- | --- | --- | --- | --- |
|  | Mean or number | SD or % | Mean or number | SD or % |  |
| Age (years) | 60.4 | 9.9 | 58.8 | 10.1 | 0.26 |
| Gender: male | 36 | 36.7 | 34 | 33 | 0.58 |
| Screening Systolic Blood Pressure (mm Hg) | 136.7 | 23.3 | 137.2 | 22.6 | 0.89 |
| Screening Diastolic Blood Pressure (mm Hg) | 74.3 | 12.9 | 77.7 | 12.6 | 0.06 |

\*RZS: random zero sphygmomanometer; SD: standard deviation

**S1 Table** shows the BP levels of participants by three different device setups: OSM, OMM and RZS and the two different measurement orders (Procedures A and B). Among all 201 participants, mean BP levels were the lowest when measured by the RZS method compared with the OSM method (difference for SBP: −0.6 mm Hg, *P=*0.15; DBP: −0.4 mm Hg, *P*=0.38) and OMM method (difference for SBP: −1.1 mm Hg, *P*<0.0001; DBP −0.8 mm Hg, *P*<0.001).

Bland and Altman plots of individual BP differences between two devices/modes (OSM/OMM/RZS) against mean BP level of the two devices/modes are presented in **Fig 2** and **S1 Fig**: (a) OSM vs. RZS, (b) OSM vs. OMM and (c) OMM vs. RZS. Most data points (>90%) fell within the limits of mean difference ± 2SD of the difference, and the agreement was substantially higher between OMM and RZS, probably given their simultaneous measurement. However, consistent biases were suggested as inverse linear relationships between the SBP difference and mean SBP level were observed between OSM and RZS methods (**Fig 2a**), and OSM and OMM methods (**Fig 2b**); and there was a positive relationship of BP values between OMM and RZS (**Fig 2c**).

**Fig 2.**
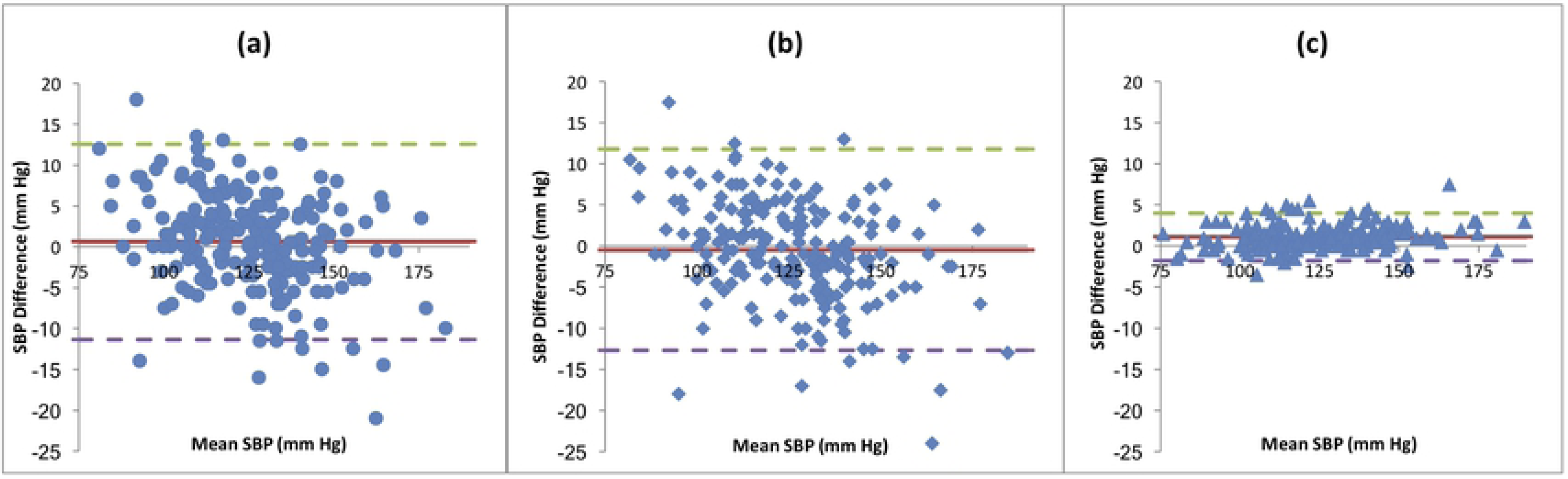
Bland-Altman Plots: individual differences plotted against means of systolic blood pressure obtained with any two different methods. (a) Omron Single Mode vs. Random Zero Sphygmomanometer; (b): Omron Single Mode vs. Omron Manual Mode; (c) Omron Manual Mode vs. Random Zero Sphygmomanometer. SBP: systolic blood pressure. Solid lines: represent overall mean difference; dashed lines: represent 95% confidence interval of overall mean difference.

With OSM the method most commonly used in large epidemiologic studies, studies of BP values measured by RZS need to be converted to OSM BP values for BP comparisons over time. Therefore, the following analyses are focused mainly on the comparisons of BP values between OSM and RZS.

Data on potential factors related to BP differences (gender, age group, BP level and device order) are presented in **Table 2**. The differences in BP values between the two devices were significantly larger among men than women (2.5 vs. −0.4 mm Hg for SBP differences, *P*=0.001; 1.9 vs. −0.5 mm Hg for DBP differences, *P*=0.005). The SBP difference was significantly smaller in the older age group (1.6 mm Hg for 40-59 years old vs. −0.5 mm Hg for 60-79 years old *P*=0.01), while their DBP difference was significantly larger than those of the younger group (−0.7 vs. 1.5 mm Hg, *P*=0.007). The SBP differences between the devices were not significantly different between the two observers (*P*=0.33), although the DBP differences were significantly different between the two observers (1.1 vs. −0.5 mm Hg, *P*=0.04). Participants first measured by OSM (Procedure A) had larger SBP differences between the devices than participants who were first measured by RZS (Procedure B) (2.0 vs. −0.7 mm Hg, *P*=0.002).

**Table 2.** Potential determinants of blood pressure differences (mm Hg) between the Omron HEM-907 Single Mode and the random zero sphygmomanometer.

| Variable | N | Systolic blood pressure difference |  | Diastolic blood pressure difference |  |
| --- | --- | --- | --- | --- | --- |
|  |  | Mean | SD | Mean | SD |
| Overall | 201 | 0.6 | 6.1 | 0.4 | 5.7 |
| Gender |  |  |  |  |  |
| Men | 70 | 2.5 | 5.4 | 1.9 | 5.8 |
| Women | 131 | -0.4 | 6.2 | -0.5 | 5.5 |
| Age group |  |  |  |  |  |
| 40-59 years old | 107 | 1.6 | 6.0 | -0.7 | 5.5 |
| 60-79 years old | 94 | -0.5 | 6.1 | 1.5 | 5.7 |
| Observer |  |  |  |  |  |
| Observer 1 | 109 | 0.2 | 5.5 | 1.1 | 5.8 |
| Observer 2 | 92 | 1.1 | 6.8 | -0.5 | 5.5 |
| Blood pressure level |  |  |  |  |  |
| Normal | 79 | 2.4 | 5.6 | -1.9 | 4.9 |
| Pre-hypertension | 81 | -0.6 | 6.0 | 1.4 | 5.4 |
| Hypertension | 41 | -0.4 | 6.6 | 2.7 | 6.3 |
| Gender and age group |  |  |  |  |  |
| Men |  |  |  |  |  |
| 40-59 years old | 32 | 3.5 | 5.5 | 1.5 | 6.3 |
| 60-79 years old | 38 | 1.7 | 5.3 | 2.2 | 5.4 |
| Women |  |  |  |  |  |
| 40-59 years old | 75 | 0.8 | 6.0 | -1.6 | 4.9 |
| 60-79 years old | 56 | -2.0 | 6.2 | 1.0 | 5.9 |
| Device order |  |  |  |  |  |
| Omron Single-RZS | 98 | 2.0 | 5.5 | 1.0 | 6.0 |
| RZS-Omron Single | 103 | -0.7 | 6.4 | -0.2 | 5.4 |
| Gender and device order |  |  |  |  |  |
| Men |  |  |  |  |  |
| Omron Single-RZS | 36 | 3.2 | 4.8 | 2.6 | 6.6 |
| RZS-Omron Single | 34 | 1.8 | 6.0 | 1.2 | 4.8 |
| Women |  |  |  |  |  |
| Omron Single-RZS | 62 | 1.3 | 5.8 | 0.1 | 5.4 |
| RZS-Omron Single | 69 | -1.9 | 6.3 | -0.9 | 5.5 |

RZS: random zero sphygmomanometer; SD: Standard deviation.

### Multivariable linear regression analyses

The results of multiple linear regression models with the RZS SBP and DBP values as the independent variables are presented in **Table 3**. These equations can be used to convert BP measurements by RZS to OSM values, which enable comparisons of BP levels and calculations of BP change over time. For example, SBP of 110.0 mm Hg for a male participant by RZS would be converted to OSM reading of 115.0 mm Hg using regression model M1 (17.4863 + 0.8555×SBP_RZS_ + 3.3796×Gender_1=male_). The predicted mean BP values based on different models developed in the BP calibration study were very similar, thus the model M1 was selected to be used to convert SBP and DBP values from RZS readings to OSM readings. Further adjustment for device order, and age could only improve the model slightly as the adjusted R-squared and explained variance were not markedly changed.

**Table 3.** Established multiple linear regression models* used to convert blood pressure (BP) readings from the random zero sphygmomanometer (RZS) to the Omron (Single Mode) and adjusted R-square for each model (N=201)

| Models | Intercept | RZS value (mmHg) | Device order | Age (years) | Gender** | Adjusted R<sup>2</sup> | Predicted mean BP (mm Hg) | Root mean square error |
| --- | --- | --- | --- | --- | --- | --- | --- | --- |
| *Systolic blood pressure* |  |  |  |  |  |  |  |  |
| M0 | 17.9050<br>(2.3864) | 0.8615<br>(0.0189) |  |  |  | 0.912 | 125.4<br>(17.5) | 5.43 |
| M1 | 17.4863<br>(2.2865) | 0.8555<br>(0.0181) |  |  | 3.3796<br>(0.7723) | 0.920 | 125.4<br>(17.6) | 5.20 |
| M2 | 18.4146<br>(2.6329) | 0.8620<br>(0.0203) |  | -0.0294<br>(0.0412) | 3.4063<br>(0.7741) | 0.920 | 125.4<br>(17.6) | 5.21 |
| M3 | 15.9052<br>(2.3301) | 0.8609<br>(0.0180) | 1.9308<br>(0.7282) |  | 3.2828<br>(0.7617) | 0.922 | 125.4<br>(17.6) | 5.12 |
| *Diastolic blood pressure* |  |  |  |  |  |  |  |  |
| M0 | 9.9934<br>(2.2948) | 0.8599<br>(0.0328) |  |  |  | 0.774 | 69.2<br>(10.1) | 5.46 |
| M1 | 9.5481<br>(2.2447) | 0.8533<br>(0.0322) |  |  | 2.5939<br>(0.7909) | 0.784 | 69.2<br>(10.2) | 5.33 |
| M2 | 1.7953<br>(3.4439) | 0.8715<br>(0.0322) |  | 0.1103<br>(0.0377) | 2.3853<br>(0.7794) | 0.792 | 69.2<br>(10.2) | 5.23 |
| M3 | 9.0022<br>(2.3709) | 0.8574<br>(0.0327) | 0.5535<br>(0.7657) |  | 2.5648<br>(0.7929) | 0.784 | 69.2<br>(10.2) | 5.34 |

Presented as beta coefficients (standard errors)
\*Multiple linear regression models were established following:
Model 0 (M0): Only include BP value by RZS;
Model 1 (M1): M0 plus adjustment for gender;
Model 2 (M2): M1 plus adjustment for age;
Model 3 (M3): M1 plus adjustment for device order of BP measurement in procedures (first by Omron Single Mode or by RZS).
\*\*1=Male, 0=Female.
BP: Blood pressure; RZS: random zero sphygmomanometer; SBP: systolic blood pressure; DBP: diastolic blood pressure.

The agreements of BP classes between the OSM and RZS, and between the OSM and calibrated RZS BP values are presented in **Table 4**. The observed percent agreement of the classification of BP was 89.1% (179/201), which was the same for both non-calibrated RZS measurement and calibrated RZS measurement. The percentages of agreement were high across all BP categories, though we found that agreement rate (79.0% for un-calibrated RZS measurement and 84.0% for calibrated RZS measurement) was relatively lowest in the pre-hypertension group. When RZS measurement was calibrated, the agreement rate was higher for the pre-hypertension group. The weighted kappa coefficient was 0.87 (95% confidence interval: 0.82, 0.92) between OSM and un-calibrated RZS, and 0.86 (95% confidence interval: 0.81, 0.92) between OSM and calibrated RZS (data for kappa coefficient not shown).

**Table 4.** Agreement of classifications of participants in categories of blood pressure levels between Omron Single Mode measurement and random zero sphygmomanometer measurement or calibrated random zero sphygmomanometer blood pressure values (N=201)

|  | Classification by Omron Single Mode measurement |  |  |  |
| --- | --- | --- | --- | --- |
|  | Normal | Pre-hypertension | Hypertension | Total |
| *Classification by RZS measurement* |  |  |  |  |
| Normal | 76 (96.2%) | 7 (8.6%) | 0 (0%) | 83 (41.3%) |
| Pre-hypertension | 3 (3.8%) | 64 (79.0%) | 2 (4.9%) | 69 (34.3%) |
| Hypertension | 0 (0%) | 10 (12.4%) | 39 (95.1%) | 49 (24.4%) |
| Total | 79 (100%) | 81 (100%) | 41 (100%) | 201(100%) |
| *Classification by calibrated RZS measurement* |  |  |  |  |
| Normal | 75 (94.9%) | 7 (8.6%) | 0 (0%) | 82 (40.8%) |
| Pre-hypertension | 4 (5.1%) | 68 (84.0%) | 5 (12.2%) | 77 (38.3%) |
| Hypertension | 0 (0%) | 6 (7.4%) | 36 (87.8%) | 42 (20.9%) |
| Total | 79 (100%) | 81 (100%) | 41 (100%) | 201(100%) |

Presented as Number (% for the column)
RZS: random zero sphygmomanometer
BP classes: normal (<120 mm Hg), pre-hypertension ( ≥ 120 mm Hg and <140 mm Hg), and hypertension ( ≥ 140 mm Hg) according to JNC 7.

## Discussion

In order to make BP values measured by different devices comparable, a BP calibration study was conducted, and the BP values from RZS, OSM and OMM were compared and equations for calibration generated. We found that the difference in BP values between the two devices related to age, gender, and BP level.

Mean BP values measured by RZS were lower than those by Omron AOD, consistent with results of previous studies [20, 21]. The Hawksley RZS has been known to underestimate SBP and DBP compared with a standard HgS [25-28]. Mackie et al.’s pooling analyses of 9 studies that compared RZS and HgS showed that RZS underestimates SBP, on average by 1.4 mm Hg and DBP by 2.0 mm Hg when compared with the standard HgS [28]. BP measurements by RZS failed the quality standards of the AAMI and the BHS, and RZS in 2001 was no longer recommended by the ESH for clinical or epidemiologic research [5]. The Omron HEM-907 has been validated previously according to the ESH, BHS and AAMI standards [11-14], and is an appropriate device for clinical practice and epidemiological studies.

The mean differences in SBP and DBP between RZS and OSM were less than 1.0 mm Hg in our study. Its carefully designed protocol and comprehensive training procedures minimized the possibility of systematic bias from observers. Moreover, to test the potential influence on BP values of different observers, an indicator variable for observer was entered into the regression models in an extra sensitivity analysis. The non-significance of the parameter estimate for this indicator variable indicates that potential variations between observers did not affect the BP differences in this study.

A German cohort study compared RZS with an Omron AOD (Omron HEM-705CP) in 2,365 participants who were randomized into groups by device order [20]. It documented mean differences between the devices (AOD-RZS) of 3.9 mm Hg for SBP and 2.6 mm Hg for DBP. We did not observe such a large difference between the two devices in our current study: only 0.6 mm Hg for SBP and 0.4 mm Hg for DBP on average. Differences in study design, and the different Omron device may account for the different results. The German cohort study also randomized participants by device orders, but there was a long time gap (22 minutes on average) between BP measurements by Omron or RZS, and there was no measurement by Omron and RZS simultaneously. In the current BP calibration study, 3 minutes rest was stipulated between measurements by devices. As BP level is highly variable with time, a longer time gap between measurements may result in increased variability of BP values by devices. Use of different Omron machines (Omron HEM-907 versus HEM-705CP) may also play an important role in the differences of the two studies, as different AODs use different BP measuring algorithms to calculate BP levels; these are not disclosed by the manufacturers.

The change trends of BP differences with BP level demonstrated by the Bland and Altman plots were inconsistent with results in the German cohort study, in which the BP difference between the Omron and RZS increased with BP level across the whole BP range; the higher the level, the larger the difference between AOD and RZS [20]. In the present study, BP differences between the two devices depended on BP level, with the differences being greater at both lower and higher BP levels. This suggests that device differences in BP measurements may be larger and have greater variability when the BP values are relatively low or high. This finding is also consistent with previous studies which documented that accuracy of AODs tends to decrease at extremes of BP due to decreased precision of the algorithms when the BP value lies outside the mid-range zone [29].

The BP differences between the two devices of the present study were greater among men than women, and BP values by AOD were slightly lower than RZS in women whereas BP values by AOD were higher in men. These results are consistent with another BP calibration study which enrolled 1,729 participants in the northern Sweden MONICA (Multinational MONItoring of trends and determinants in CArdiovascular disease, MONICA) population survey comparing RZS with Omron M7 AOD[21]. We also found that the DBP difference increased with age independent of gender in our study. The differences in BP values by gender and age here might be due to differences in white coat effects [30-33], as some previous studies reported that men and women had different white coat BP responses [30, 33]. The current study documented a mean SBP difference between OSM and RZS of 2.5 mm Hg for men and −0.4 mm Hg for women. Previous studies have shown that women have larger white-coat effects than men [30-32]. It has been shown that the use of AOD reduces the white-coat effect compared with manual BP measurements by HgS or RZS [34]. Therefore, a larger BP increase in women than in men due to white-coat effects was expected when BP was measured manually by RZS, while BP increase due to white-coat effects might be minimized when BP was measured by automated OSM. In this case, a smaller BP difference between OSM and RZS (BP_OSM_-BP_RZS_) would be expected in women, as seen in the current study.

Agreement between OMM and RZS was high in the current study, compared with the agreement between OSM and RZS. SDs of the BP difference by the two devices (RZS and OMM) are very small, which is consistent with the BP calibration study in the CARDIA Study. The CARDIA Study also did a similar BP calibration study which compared the Omron HEM-907 and RZS [35], but it only included simultaneous measurements by RZS and OMM and generated an equation to calibrate BP values. As shown by the results from the current study, there might be systematic bias between OSM and OMM as well. If the calibration study used the manual mode, while the data collection for the prospective study used the single mode, the results of BP change could still be biased. The current study design enabled all of the comparisons between RZS, OSM and OMM, and can be used to reduce systematic biases of BP recordings by devices/modes in population based studies.

The observation of differences between the RZS and the Omron machine implies that misclassification of hypertensive and normotensive persons and values of BP changes might be biased if different devices are used. It also suggests that the BP range of study participants might influence the validity of equations calculated for calibration. Development of calibrated equations for specific BP ranges might therefore need to be considered.

Moreover, an order effect on SBP has been documented in our study as well, similar to previous studies [20, 21]. One potential explanation is observer bias. Once the observer of the RZS measurement knew the BP level obtained from the Omron device, he or she might be able to more accurately detect the phase I (SBP) and phase V (DBP) Korotkoff sounds [20]. However, as measurements of Omron and RZS in the current study were obtained by two independent observers in separate seats without any communication during the whole process (‘blinding to BP readings measured by each other’), and the random change in the zero level of the RZS was applied, this explanation is not very convincing. Another explanation is that the first reading is typically the highest when a series of BP readings is taken [22]. For this reason, we discarded the first reading by each device, similar to other studies. Moreover, we applied a randomization procedure of device order in the study design and included both procedures in our regression analyses of the complete data set. Therefore, the influence of device order should have been minimized.

Our model obtained for calibration is simple, as the variables of gender are easily obtained. Even though no previous study compared RZS with Omron HEM 907 (Single Mode) in different ethnic groups, the parameter estimates of the equation in the CARDIA BP calibration study are very close to the parameter estimates in the current study comparing RZS with Omron HEM-907 (Manual Mode). Therefore, it can be suggested that this equation comparing RZS with Omron Single Mode could be applicable to other populations.

In a summary, several variables, including age and gender, contribute to the BP differences between the RZS and AODs. Equations were developed taking those variables into account to convert BP values by RZS to Omron AOD. These equations can be used to compare BP values between studies or a shift of the BP distributions over time using different devices for BP measurements

## Acknowledgements

We thank Dr Jeremiah Stamler for his critical and helpful comments on final draft of the manuscript.

## Supporting information

**S1 Appendix. Converting random-zero to automated oscillometric blood pressure values using Deming Regression.**

**S1 Table. Mean blood pressure levels of participants and mean differences of blood pressure values by Omron single mode, Omron manual mode and random-zero sphygmomanometer for all participants (N=201) and stratified by procedures.**

**S2 Table. Results of Deming Regression models for converting blood pressure (BP) readings from the random zero sphygmomanometer (RZS) to the Omron (Single Mode).**

**S3 Table. Agreement of classifications of participants in categories of blood pressure levels between Omron Single Mode measurement and calibrated random zero sphygmomanometer blood pressure values.**

**S1 Fig. Bland-Altman Plots: individual differences plotted against means of diastolic blood pressure (DBP) values obtained with any two different blood pressure methods (devices/modes) among 201 study participants in the blood pressure calibration study.** (a) Omron Single Mode vs. Random Zero Sphygmomanometer; (b): Omron Single Mode vs. Omron Manual Mode; (c) Omron Manual Mode vs. Random Zero Sphygmomanometer. Solid lines: represent overall mean difference; dashed lines: represent 95% confidence interval of overall mean difference.

